# Virtual Tissue Expression Analysis

**DOI:** 10.1101/2023.11.16.567357

**Authors:** Jakob Simeth, Paul Hüttl, Marian Schön, Zahra Nozari, Michael Huttner, Tobias Schmidt, Michael Altenbuchinger, Rainer Spang

**Affiliations:** LIT – Leibniz Institute for Immunotherapy, c/o Universitätsklinikum Regensburg, Franz-Josef-Strauß-Allee 11, 93053 Regensburg, Germany; Statistical Bioinformatics, Faculty of Informatics and Data Science, University of Regensburg, Am Biopark 9, 93053, Regensburg Germany; Department of Medical Bioinformatics, University Medical Center Göttingen, Goldschmidtstr. 1, 37077 Göttingen, Germany

**Keywords:** cell-type specific expression, cell-specific gene regulation, cellular composition, deconvolution

## Abstract

**Motivation:** Bulk RNA expression data is widely accessible, whereas single-cell data is relatively scarce in comparison. However, single-cell data offers profound insights into the cellular composition of tissues and cell-type-specific gene regulation, both of which remain hidden in bulk expression analysis.

**Results:** Here, we present tissueResolver an algorithm designed to extract single-cell type information from bulk data, enabling us to attribute expression changes to individual cell types. The outcome is a virtual tissue that can be analyzed in a manner similar to single-cell RNA-seq data. When validated on simulated data tissueResolver outperforms competing methods. Additionally, our study demonstrates that tissueResolver reveals previously overlooked celltype specific regulatory distinctions between the activated B-cell-like (ABC) and germinal center B-cell-like (GCB) subtypes of diffuse large B-cell lymphomas (DLBCL).

**Availability and Implementation:** R package available at https://github.com/spang-lab/tissueResolver. Code for reproducing the results of this paper is available at https://github.com/spang-lab/tissueResolver-docs.

## INTRODUCTION

Present day single cell studies describe the biology of tissues in unprecedented detail. They molecularly characterize many thousands of cells and unveil the extensive spectrum of cellular phenotypes present within a tissue.

When comparing two categories of tissues, single-cell data directly associates differentially expressed genes with their specific cell origins. For instance, if a group of mitotic genes shows upregulation at the tissue level in a tumor, single-cell data can distinguish whether these genes are expressed in actively dividing tumor cells or, conversely, in expanding immune cell populations that are combating the tumor.

However, scRNA-seq data is expensive to generate and datasets are often limited in the number of independent samples, focusing on the within-sample heterogeneity rather than on large sample populations. Furthermore, due to the relatively recent development of this technology, clinical follow-up data for patients are scarcely available, if at all. In contrast, bulk expression data has been systematically gathered in substantial quantities and is well-annotated from a clinical perspective. These data are readily accessible through sources such as the Cancer Genome Atlas (TCGA) [1] or the International Cancer Genome Consortium (ICGC) [2].

The quantification of cell populations from bulk tissue sequencing data, known as tissue deconvolution, is a vibrant area of research [3] and deconvolution tools provided important insights into the nature of tissues [4]. Their effectiveness stems from their ability to harness large bulk datasets.

More recent devonvolution methods distinguish themselves by learning the distinctive characteristics of the specific cell types of interest from single cell data [5, 6]. The single cell data is used to construct reference profiles for each cell type. While this approach is beneficial for estimating cell type frequencies, it unavoidably simplifies the intricate regulation processes within individual cell populations, which cannot be fully encompassed by a cell type label alone.

Recent methodologies fine-tune the reference profiles to the particular tissue type of interest [6, 7, 8, 9]. However, even with these advancements, the within-tissue diversity of cells still remains concealed.

The challenge, therefore, lies in bridging the gap between these two available sources: bulk RNA sequencing data from extensive cohorts with comprehensive clinical information and single-cell RNA sequencing, which offers a precise depiction of tissue characteristics.

When we have knowledge of cell type frequencies within the bulk tissue, it becomes feasible to attribute bulk gene expression of specific genes to individual cell types [10, 11, 12, 9].

Nonetheless, a strict cell type counting approach fails to encompass the considerably more nuanced, gradual molecular variations that occur within cells of the same type. Cells can undergo activation or reprogramming in numerous different ways, and their continually evolving molecular phenotypes give rise to intricate trajectories in a multidimensional space, which is far more intricate than a mere list of discrete cell types [13].

In groundbreaking work, BayesPrism [14] departs from the notion of fixed cell types and instead strives for a simultaneous estimation of cellular phenotypes and their frequencies within a tissue. BayesPrism is an exceptionally comprehensive full Bayesian model that quantitatively encompasses a wide array of facets in tissue biology, spanning from cellular composition to gene regulation, and from cell plasticity to regulatory dynamics. This multifaceted model comprises two primary steps: In a first deconvolution step, the method constructs a prior based on cell type-specific expression profiles obtained from single-cell RNA-seq data. It uses this prior to jointly estimate the posterior distribution of both the cell type composition and the cell type-specific gene expression profiles from bulk RNA-seq data of tumor (or non-tumor) samples. In a subsequent embedding step the Expectation-Maximization (EM) algorithm is used to approximate the tissue’s gene expression profiles. It does this by modeling it as a linear combination of gene programs while conditioning on the inferred expression and the cell fractions estimated by the deconvolution module. Similarly, ISLET [15] is another probabilistic approach that is promising.

Here, we introduce “tissueResolver”, an alternative strictly deconvolution based approach for generating virtual tissues. Unlike standard deconvolution approaches, this method operates without the need for predefined cell type labels. Instead, it autonomously identifies distinct cell populations exhibiting regulatory differences across various tissues.

We conducted performance tests on tissueResolver using simulations involving bulk data derived from modified single-cell data. In this process, we also explored the boundaries of its applicability. We observed that tissueResolver excels over BayesPrism when applied to the same scenarios, particularly in terms of its ability to provide higher resolution and specificity when attributing expression changes to specific cell types.

In a case study involving diffuse large B-cell lymphomas (DLBCL), tissueResolver provided fresh insights into the subtle expression differences between its activated B-cell-like (ABC) and germinal center B-cell-like (GCB) subtypes. Notably, these differences extended beyond the tumor cells themselves to encompass various elements of the tumor microenvironment. Specifically, we were able to attribute some changes to a small cluster of cells exhibiting expression profiles resembling fibroblasts.

## ALGORITHM

### Computation of virtual tissues

We approach the following problem: Given a bulk expression profile and a library of single cell profiles, we aim to determine the optimal subset of cells and their corresponding weights in such a way that the sum of their weighted expressions closely approximates the bulk expression profile in euclidean distance.

Formally, we minimize:

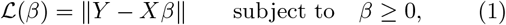

where 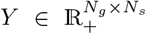 is a matrix of *N*_*s*_ samples, 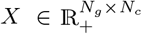 the library of single cell profiles, and *N*_*g*_ is the number of genes in the profiles. The parameters that we wish to determine are the entries of the weight matrix 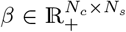 . We do so using non-negative regression [16] resulting in a sparse non-negative estimate of *β*. To further stabilize the estimate, we bootstrap the single cell data *N*_*b*_ times using a random 10 % percent of cells and rerun the regression analysis. This results in a set of *N*_*b*_ different single cell libraries and corresponding sparse weight estimates that are analysed separately and in the end results are averaged.

When following this approach, we assume that the single cell library is comprehensive and contains cells of most types and states present in the bulk tissue. We further assume that the bulk expression profile can be effectively reconstructed by selecting the appropriate cells from this extensive collection. However, not all cell types or cell states from the library will be present in every bulk tissue. Therefore, we use a sparse estimate of *β*. Our approach is like picking out specific Lego tiles from a vast collection to assemble them in a way that best fits a given assembly, in this case the bulk expression profile.

The resulting combination of weights *β* and single cell profiles *X* essentially decomposes the bulk tissue into single cell profiles. Instead of having a single expression value for a gene, we now possess thousands of values, each corresponding to an included cell. Consequently, we refer to these sets of values as a “virtual tissue”. The virtual single cell profiles can be further subjected to downstream analyses in a manner similar to real single cell data.

It is worth noting that this approach resembles a deconvolution approach in which each single cell is treated as its own “cell type”. In the collection, there may be many cells of the same type, like CD8+ T-cells, but they may be in different expression states, such as activated and inactivated. If the algorithm predominantly selects activated T-cells in tissue A and inactivated T-cells in tissue B, we can observe changes in gene expression that are clearly associated only to T-cells. Changes in gene expression were already observable in the bulk profile. However, it was only after the deconstruction of the bulk into single cells that it became clear that these genes were differentially expressed in CD8+ T-cells.

In classical deconvolution approaches, the matrix *X* contains only a few columns, typically one for each included cell type. In this setup, the linear equation system *Y* − *Xβ* is over-determined. However, in tissueResolver, *X* includes many more columns, one for every cell in the library, which necessitates regularization techniques to handle the increased dimensionality.

We chose non-negative regression for learning the weights *β*. This approach not only ensures that the weights remain non-negative, which is crucial for interpretability, but it also leads to a sparse and regularized estimate of *β* [16] without additional regularization parameters, and avoids that similar cells can cancel each other out. Cells that are not similar to cells present in the bulk will be assigned weights that are very close to zero. In contrast, for redundant cells that contribute much of the expression found in the bulk, a few representative cells will be chosen that will be assigned large weights.

Selecting an appropriate library is important. The number of non-zero weights tends to be higher when there is a significant overlap between the bulk tissue and the single cell library. This overlap is more likely when bulk and single cell data are derived from similar tissues. If cell identities are missing in the library, however, certain genes will not be represented well in the virtual tissue, *Xβ*, and we find large residuals for these particular genes.

Bootstrapping further helps in quantifying such uncertainties. Large residuals of the final model as well as unstable bootstrap results point to problems in the choice of the library. Furthermore, the gene-wise variance across bootstrap samples serves as a valuable quality metric, assessing the extent to which a particular gene is effectively represented in the single cell dataset. Only genes that are well covered will be interpreted in subsequent analysis.

### Cell frequencies and cell-type specific gene expression from virtual tissues

The previous section did not require any cell type labels for the cells in the library. In fact, tissueResolver operates without requiring cell type labels, distinguishing it from other tools like BayesPrism, see also “Simulations”. However, the introduction of cell types can be advantageous for enhancing our comprehension of tissue biology, as it situates virtual tissues within the framework of conventional histology and cell biology. In the following, we assume a set ℳ of *N*_*L*_ distinct cell-type labels and denote with 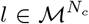 the vector labeling every single cell in *X*. Note that there are many complementary ways to label cells, e.g. cell type annotation, cluster membership, or activation and cell cycle states, and for every choice of *l*, the same virtual tissue can be re-analysed.

For every cell label *a* ∈ ℳ, we can compute effective cell-frequencies by summing over the respective weights,

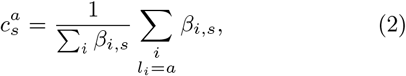

which we repeat for every bootstrap sample in order to compute bootstrap averages and errors in the end. This yields the effective frequency 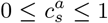 of cells of type *a* in sample *s*.

Note that the values of 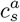 can only be meaningfully interpreted when comparing the frequency of the same cell type across different tissues. Comparing different cell types within the same tissue, on the other hand, is not valid due to variations in the total amount of expressed RNA across cell types.

Estimating cell type-specific gene expression follows a similar approach. To calculate the estimated expression profile of a cell type *a* in sample *s*, we sum the explained expression over all cells of that particular type,

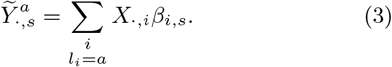

Note that the sum over all cell labels recovers the estimated total gene expression, 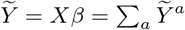.

Both cell frequencies and cell type specific expression profiles shape the bulk profile. TissueResolver disentangles these contributions, allowing for a more comprehensive understanding of the underlying tissue heterogeneity. For instance, when we aim to examine differences in cell type-specific gene expression across two classes of tissues, it is essential to account for variations in the abundance of these cells in each tissue. To achieve this, the cell type-specific expression is straightforwardly normalized by its relative weight, 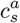,

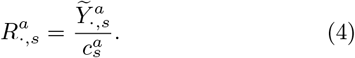

### Quality scores and gene selection

It is crucial to thoroughly assess virtual tissues before drawing any conclusions based on their results. Some genes within the virtual tissue may not align well with the actual bulk tissue due to the absence of certain cell types in the single cell dataset or due to technological disparities between bulk and single cell sequencing. Additionally, individual virtual tissues may have failed to fully account for their respective bulk profiles. Once more, this could indicate inadequate single cell libraries lacking essential cell identities required for accurate representation of the bulk samples. In the following section, we will explore how spurious genes and virtual tissues can be identified and subsequently excluded from further downstream analysis.

On every bootstrap sample *k*, we compute for every gene *g* and sample *s* the relative residual,

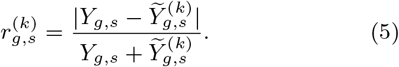

Note that 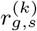 takes on values in the range [0, 1], with values close to zero indicating a perfect fit of gene *g* in sample *s*, while a value near one indicates that the explained gene expression substantially deviates from the actual gene expression in the bulk.

By averaging 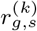 over bulk samples and bootstrap samples, we can assess how effectively gene *g* is reconstructed across all virtual tissues. This results in the gene-specific quality score 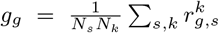 . In contrast, averaging over all genes of a virtual tissue, 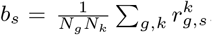, gives a quality score for individual virtual tissues.

To judge the robustness of the fit, it is also useful to consider mean bootstrap variances of the above quantities, e.g., 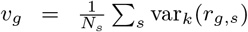. Very large values (with respect to its mean value) indicate large fluctuations in the gene expression of the individual bootstrap samples that result from a sensitivity to small communities of individual, possibly outlier cells. A detailed discussion of quality scores based on the case study shown below can be found in the supplemental material.

## SIMULATIONS

To evaluate the performance of tissueResolver and to compare it to BayesPrism [14], we introduce artificial cell-type specific gene expression changes to a single cell DLBCL dataset [17]^1^. We only change one cell type, effectively altering cell type specific gene regulation, and challenge tissueResolver to identify (a) the modified genes and (b) to attribute the changes to only the modified cell type.

The data [17] comprised 28,416 single cells obtained from eight lymphoma patients, with a total of 6,639 CD8+ T cells — the cell type we modified. We randomly divided the eight patients into two groups of four each. The cells from the first group were utilized to create libraries *X*, while those from the second group were employed to generate artificial bulk profiles.

We randomly selected half of the CD8+ T cells in both patient sets and multiplied the expression of a varying number of randomly selected genes |𝒢_mod_| by different factors ranging from 2^0.5^ ≈ 1.41 to 2^2^ = 4. The modified genes were not exclusively expressed in CD8+ T cells, but only the fraction that originated from CD8+ T cells was modified, for details see supplemental section “Simulation details”.

From the second set of samples, we created 100 artificial bulk profiles by randomly selecting 500 single cells and summing over their profiles. Out of these 100 artificial bulks, 50 were drawn from the set where CD8+ T cells were modified, forming class I. The remaining 50 samples exclusively included unmodified T cell profiles, constituting class II. By construction, class I and II only exhibited differences in gene expression within the CD8+ T cell compartment. Expression profiles in all other cell types, as well as the cellular composition, remained identical in both groups.

We run both tissueResolver and BayesPrism to estimate cell-type specific gene expression for all cell types. Both algorithms were provided with the cells from the first set as reference data and the 100 bulk profiles generated from the independent second set of samples as bulk inputs. We assessed the performance of both methods in (a) identifying the modified genes and (b) attributing the differential expression exclusively to CD8+ T cells. Separately for each cell type, we ranked genes based on differential gene expression and computed receiver operating characteristics (ROC) curves to identify the modified genes (see Figure 2a). From these ROC curves, we calculated the areas under the curve (AUC) for various log-fold changes and different numbers of manipulated genes.

**Fig. 1.**
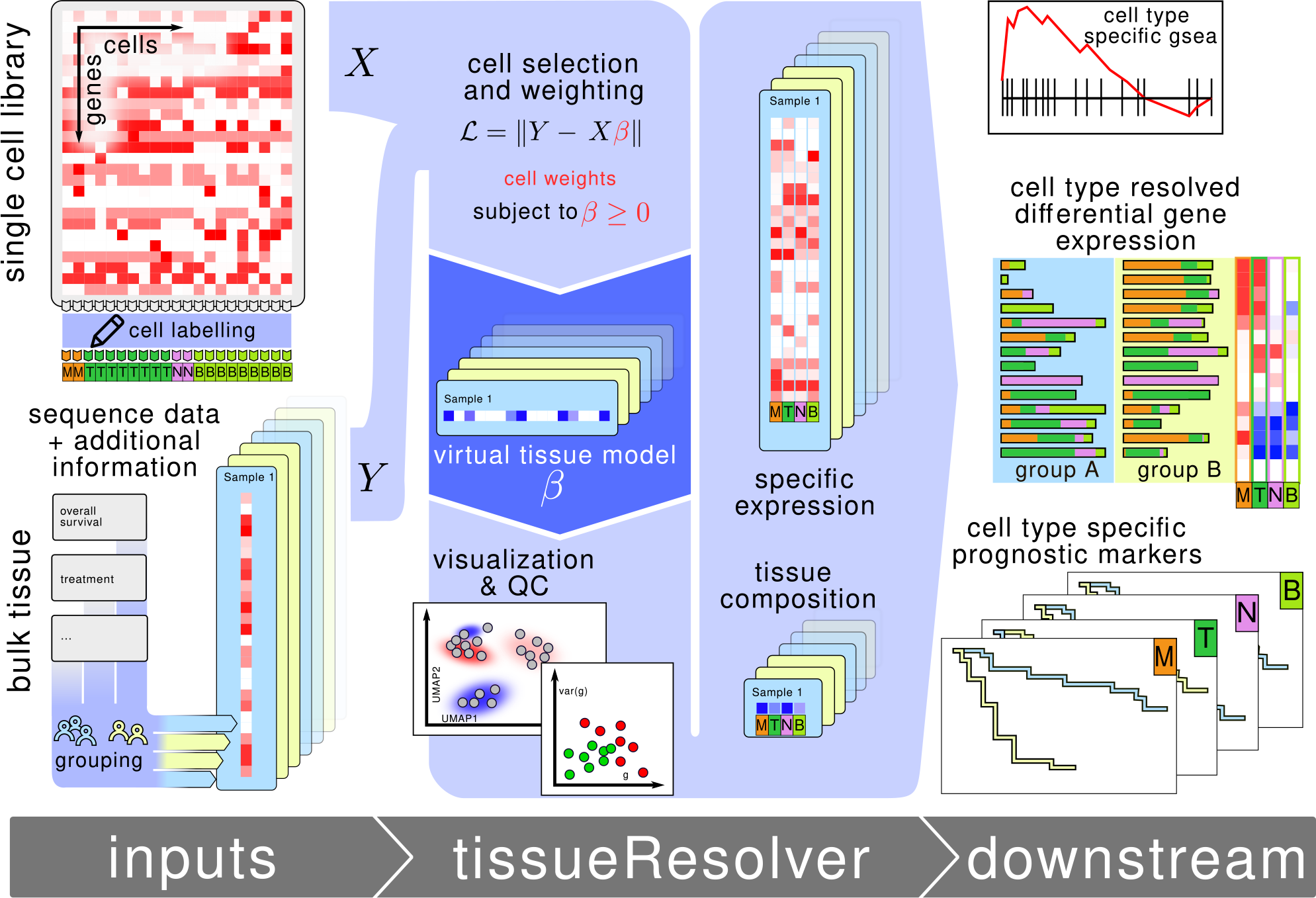
Schematic representation of the tissueResolver workflow: The application of tissueResolver requires two input datasets: Bulk tissue profiles (*Y*) and single-cell data (*X*). TissueResolver assigns positive weights (*β*) to the single-cell profiles, attempting to adjust the combination *Xβ* to closely match the bulk expression Y. The resulting model is referred to as “Virtual Tissue” and allows for a detailed analysis similar to the approach used with complete single-cell datasets. In this context, different-sized cell populations are represented by larger weights of representative cells. The obtained Virtual Tissues can be initially analyzed through the visualization of selected cells along with their weights in low-dimensional embeddings in a familiar manner. For further analyses, defined cell populations (cell types, cell states, or cell clusters) can be utilized. By using these labels, tissue compositions can be calculated, and their contribution to expression differences can be distinguished from those arising due to gene regulation in specific cell populations. In principle, these analysis steps can be performed individually for each sample. However, in many studies, bulk data is assigned to groups (tumor type, survival, treatment response). The full power of tissueResolver becomes particularly evident when these groupings are considered, allowing for cell-type-specific differential gene expression analysis between these groups.

**Fig. 2.**
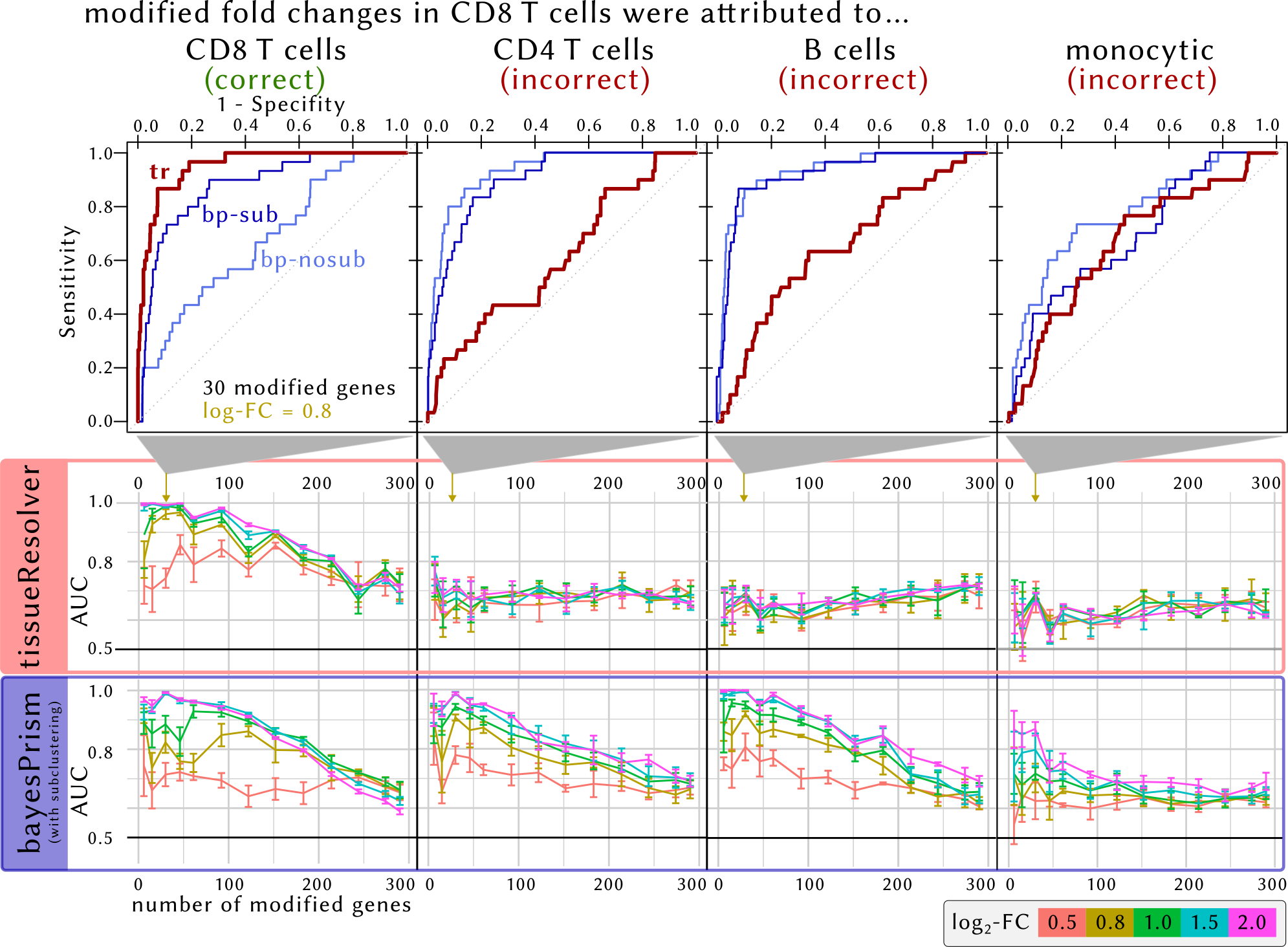
Results of simulations performed for the evaluation of tissueResolver and BayesPrism. Top row: Receiver operating characteristics for the modified CD8 T-cells and the (unmodified) CD4 T, B, and monocytic cells, computed for finding the 30 modified genes in the top differential expressed genes of the respective cell type for a change in foldchange of 2^0.8^ ≈ 1.74. Two variants of BayesPrism were run: One in which only the annotated cell type was provided (“bp-nosub”) and one in which each annotated cell type was sub-clustered further (“bp-sub”), see supplement for details. Second and third rows: Extracted area under the curve (AUC) values of tissueResolver (red) and BayesPrism after subclustering of every annotated cell type (blue) for finding a modified gene in the top differential genes of the given cell type as a function of the number of modified genes and the fold changes. The horizontal black line indicates an AUC of 0.5 and represents the null hypothesis. Values significantly larger than 0.5 mean that the modified genes have been attributed to the corresponding cell type, which we expect only for the modified CD8 T-cells, whereas other cell types are expected to give values around 0.5.

We observed that both tissueResolver and BayesPrism excel in identifying the genes that exhibited differential expression in CD8+ T cells, as evident from the overall steep ROC curves. However, tissueResolver demonstrates the capability to attribute differential expression solely to the CD8+ T-cell compartment, with minimal spill-over to similar cell types like CD4+ T-cells or B-cells. In the case of tissueResolver, only the ROC curve for the CD8+ T-cell compartment displayed a steep incline, while for other cell types, the curves closely followed the identity line. The unwanted spill-over was considerably more pronounced in BayesPrism. This method identified differential expression of the modified genes across all lymphocyte cell types, whereas tissueResolver more effectively restricted this differential expression attribution to the specific CD8+ T-cell compartment. As we increased the artificial fold change, leading to more significant expression changes, it became easier for the algorithms to accurately classify the signals, even with a moderate number of selected genes. Interestingly, the increase in fold change also led to a more distinct “misclassification” of BayesPrism for the incorrect cell types CD4 T and B, improving its specificity. Both algorithms struggle when a very high number of genes is modified: In this case, the modified cell type moves closer to the unmodified, but seems to occur with a higher frequency since all relevant genes have been upregulated and it becomes impossible to single out the genes that were modified.

It’s essential to recognize that tissueResolver and BayesPrism use cell labels in different ways. BayesPrism uses them in the pre-modeling phase, where it employs a set of pre-defined cell states, which can be considered very fine-grained cell types. In contrast, tissueResolver treats each single cell profile as an independent reference profile. It is only after the method selects the most suitable cells to explain a tissue that these cells are clustered, and marker genes are utilized to assign them to cell types or cell states. In practice, these post-modeling analysis options can prove advantageous, as we will demonstrate in the case study on the lymphoma micro-environment in the next section.

## THE MICRO-ENVIRONMENT OF DIFFUSE LARGE B-CELL LYMPHOMAS

In this section, we showcase how tissueResolver explores the gene expression characteristics of tumor tissues. Our case study centers on unveiling differences in the micro-environment of two subtypes of diffuse large B-cell lymphomas (DLBCL), which ressemble different stages of B-cell differentiation: the ABC (activated B-cell like) and GCB (germinal center B-cell like) subtypes, see [18]. We analyzed 481 RNA-seq bulk samples from Schmitz et al. [19]^2^. Among these samples, 138 belonged to the GCB subtype, and 243 belonged to the ABC subtype. The single-cell library incorporated the combined data from Roider et al. [20]^3^ and Steen et al. [17], without further data harmonisation steps, allowing tissueResolver to determine the best-fitting profiles. This dataset comprised a total of 63,700 labeled single cells from 20 patients, covering ABC and GCB DLBCL, as well as follicular lymphomas (FL) and benign reactive lymph nodes (rLN), see supplemental Table 1 and fig. 3 for further details

**Fig. 3.**
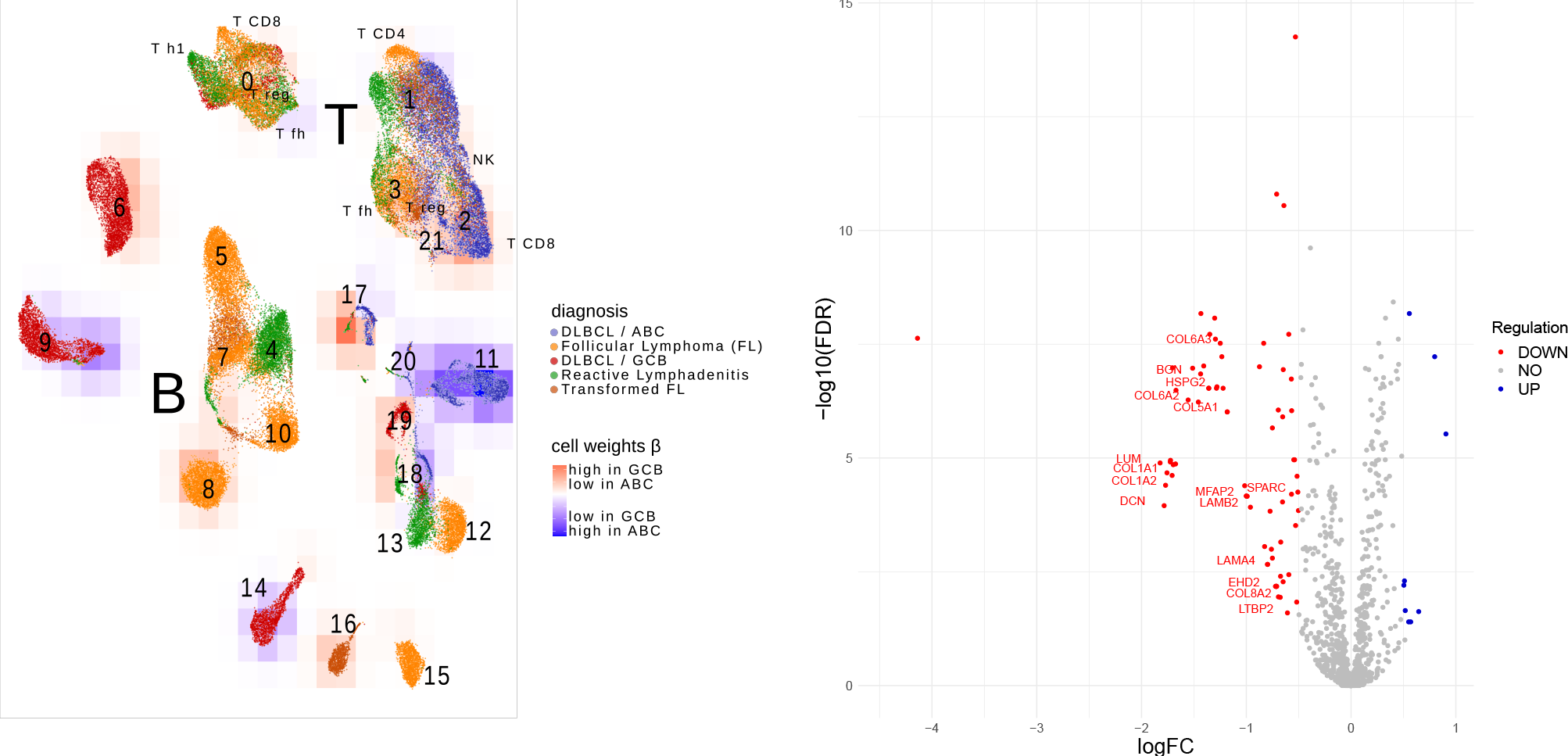
UMAP representation of the single-cell library, colors of the cells indicate the tissue of origin. The cells were incorporated into the virtual tissues of both ABC and GCB lymphomas. A red background indicates that the cells received higher weights in virtual GCB tissues, while blue indicates a preference of the cells in virtual ABC tissues.

First, we selected the 1000 most variable genes across both bulk and single-cell data (see supplemental section “Gene filtering”). Subsequently, we ran tissueResolver with 50 bootstrap samples, each consisting of 10% of the cells in the library. On average, virtual tissues comprised 148 cells (standard deviation *σ* = 51) of the 6,370 cells available in each bootstrap run. In summary, the quality control, as assessed by the scores in “Quality scores and gene selection”, indicated that virtual tissues provided a robust explanation for all bulk samples across the majority of genes. A detailed analysis of the quality control is presented in supplemental section “Quality control of virtual tissues”.

We re-clustered the combined single cells, see supplemental section “Clustering of single cells”. Figure 3 visualizes the resulting clusters in a UMAP embedding [21], while supplementary Table 2 provides additional information on the composition of the clusters.

Single cells were incorporated into both virtual ABC and GCB tissues, each assigned with distinct weights. We compared these weights and represented them using background coloring in Figure 3. A red background signifies that the cells received higher weights in virtual GCB tissues, while blue indicates a preference for the cells in virtual ABC tissues. For example, Cluster 11 contains cells that were better suited to explain ABC bulk profiles compared to GCB profiles.

We noticed large cell clusters representing tumor cells that were more prevalently integrated into one tumor subtype compared to the other. Encouragingly, virtual ABC tissues primarily included lymphoma cells from ABC single-cell data, while GCB included cells from GCB and, to a lesser extent, from follicular lymphoma single-cell data. The latter observation suggests that while follicular lymphomas have expression characteristics different from all DLBCL due to their distinct cellular composition, the malignant cell compartment alone shares similarities with GCB-DLBCL.

We also identified small areas within individual clusters showing differential weights between ABC and GCB, indicating nuanced regulatory shifts within sub-populations of a cell type. For instance, in cluster 18 (labeled as non-malignant B-cells), some cells increased in weight, while others in the same cluster decreased. Interestingly, this cluster combines cells from two patients—one with DLBCL and one with tonsillitis (T2), as indicated in supplemental Table 1 and fig 3 where we included this patient in the reactive lymphadenitis group. The fact that these cells cluster together supports the annotation as benign B-cells. However, the observation that only a subset of these cells was suitable for explaining DLBCL bulks suggests that these micro-environmental cells are nevertheless reprogrammed in lymphomas.

In clusters 0 and 17, tissueResolver also detected cellular heterogeneity by assigning differential weights within the same cluster. Originally labeled as “different T-cells” and “monocytes/myeloids”, respectively, these clusters are further examples that suggest that even when cell profiles are globally similar and cluster together, tissueResolver can identify differential regulatory effects across bulk tissues within them.

We proceeded by computing cell type specific gene expression profiles for all cell types/cell clusters.

The ABC/GCB signature was crafted to reflect distinct cells of origin for malignant lymphoma cells [18, 22, 23]. Consequently, we would anticipate that their expression differences are driven by the tumor cell compartment alone. Interestingly, these genes are not exclusively expressed in malignant B-cells [20, 17]. Therefore, we employed tissueResolver to untangle the contribution of cell clusters to the overall expression of these signature genes.

Figure 6 presents the outcomes of this analysis. The bar plots on the left validate that the signature genes are primarily but not exclusively expressed in tumor cells, with some expression observed in various non-malignant cells. The heatmap on the right attributes the observed expression differences between the two DLBCL subtypes primarily to the malignant B-cell clusters, consistent with expectations.

Several notable exceptions were observed. Firstly, the key regulator IRF4 is up-regulated in ABC, not only in the lymphoma cells but also in non-malignant plasmablasts (cluster 20), aligning with its recognized physiological role in B-cell fate determination [24]. Secondly, three genes in the signature, SPINK2, KRT8, and HLA-DAQ1, surprisingly showed expression changes attributed to non-malignant cell compartments. While these genes exhibited substantial expression contributions from malignant cells, the observed expression changes between ABC and GCB were not solely attributed to these cells but also to cells from the micro-environment. Specifically, for KRT8 and HLA-DAQ1, changes were attributed to cluster 17, which is analyzed in more depth at the and of this section and in the supplement, and which plays an important role in the explanation of a stromal signature below.

We proceeded to examine gene expression in the tumor micro-environment by dissecting the expression of stromal-1 and 2 signature genes, as outlined in [25]. These signatures were specifically crafted to complement the ABC/GCB signature, focusing on distinctions within the lymphoma micro-environment rather than the tumor cells themselves.

The stromal-1 signature has demonstrated high prognostic value. According to Lenz et al. [25], elevated values of this signature are associated with increased extracellular matrix deposition and infiltration of the tumor by histocytic immune cells, such as monocytes or macrophages. In essence, the signature is attributed solely to aspects of the cellular composition of the lymphoma micro-environment. As we will demonstrate, tissueResolver can provide additional insights into the signature, particularly concerning differential gene regulation in cells of cluster 17.

Unlike Lenz et al. we analyze these genes in the context of an ABC/GCB comparison. Many of these genes are in fact differentially expressed between these two subtypes of DLBCL.

Figure 7 provides an overview of the outcomes. The majority of signature genes showed up-regulation in GCB, and tissueResolver predominantly associated this regulation with cell cluster 17, as indicated by the green bars on the left and the prominent red color in the column associated with cluster 17 on the right. Cluster 17 is characterized by high expression of genes in the GCB subtype that are typically found in fibroblast-type cell populations.

In the single-cell data, cluster 17 contained a notably small number of cells compared to the quantities of malignant and non-malignant B-cells (Table 1). However, despite their limited count, these cells consistently appeared in the virtual tissues, indicating their significance in explaining bulk expressions. This significance is further evidenced in the relative cell weights depicted in Figure 4, where we computed the ratio of mean weights in ABC vs GCB samples for every cell cluster separately. These weights refelct the frequency of cluster 17 cells in ABC and GCB lymphomas.

**Table 1.**
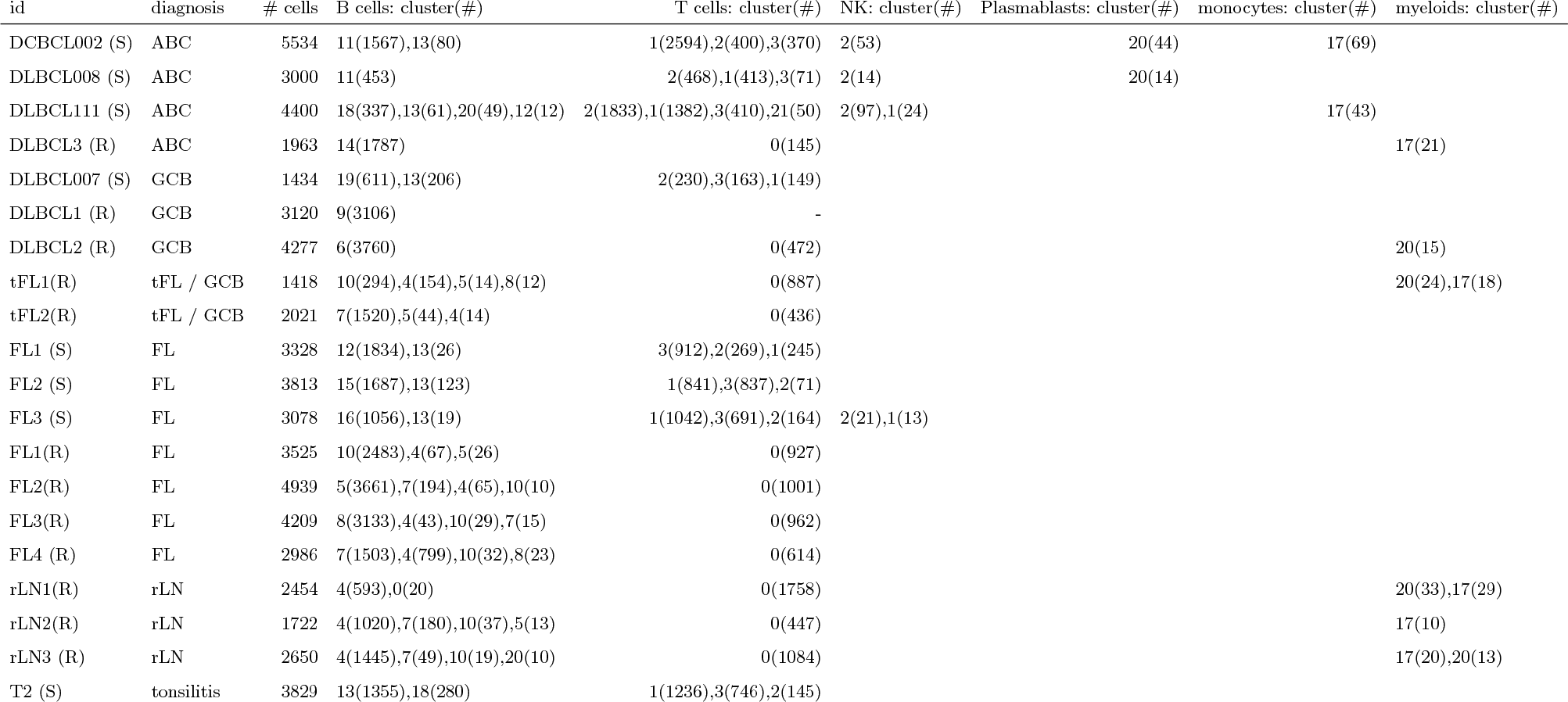
Sample identifier of scRNA-seq data of Steen et al. [17] (S) and Roider et al. [20] (R), along with their diagnoses and numbers of cells in each of our clusters. Cluster frequencies below 10 were omitted for brevity. Clusters were assigned to coarse unified labels from the original publications by majority vote.

**Table 2.**
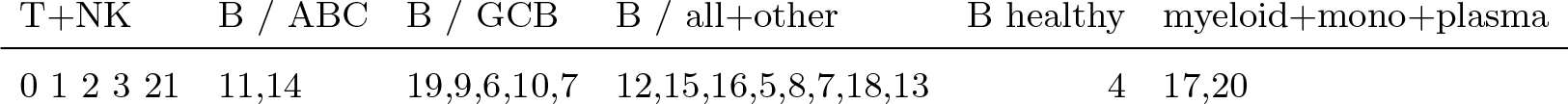
Correspondence of cell types to clusters.

| T+NK | B / ABC | B / GCB | B / all+other | B healthy | myeloid+mono+plasma |
| --- | --- | --- | --- | --- | --- |
| 0 1 2 3 21 | 11,14 | 19,9,6,10,7 | 12,15,16,5,8,7,18,13 | 4 | 17,20 |

**Fig. 4.**
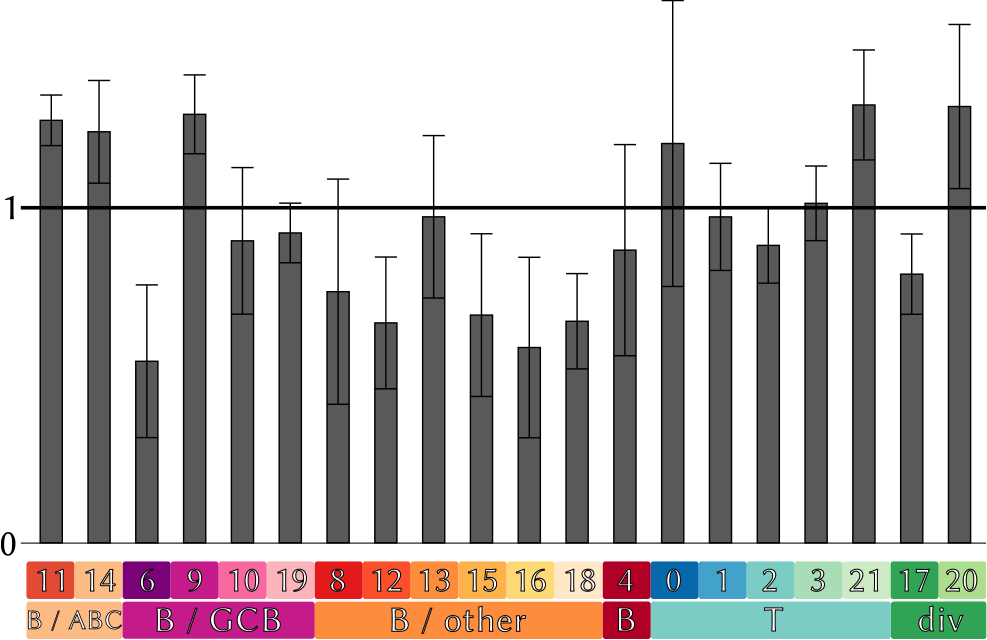
Estimated cellular composition of ABC vs. GCB tissues. Shown are ratios of average cluster weights between the two DLBC subtypes.

While we did note slightly reduced weights in the ABC subgroup, these weights alone failed to adequately explain the significant expression variations observed in the stromal genes (Figure 7). Interestingly, this cluster exhibited a substructure: a subset of its cells was predominantly associated with the virtual tissues of ABC tissues, while another subset within the same cluster was predominantly found in GCB tissues. Cluster 17 includes cells in different states, some of which were characteristic for ABC tissues while others were predominantly found in GCB tissues. This effect becomes even more apparent when we visualize the differential gene regulation within cluster 17 using a volcano plot (Figure 5) comparing ABC and GCB gene expression only in cells of cluster 17. Gene labels in Figure 5 highlight stromal signature genes, suggesting that if cluster 17 cells were utilized to define a stromal signature, the resulting gene list would closely resemble the genes identified by [25]. Further details are available in the supplement.

**Fig. 5.**
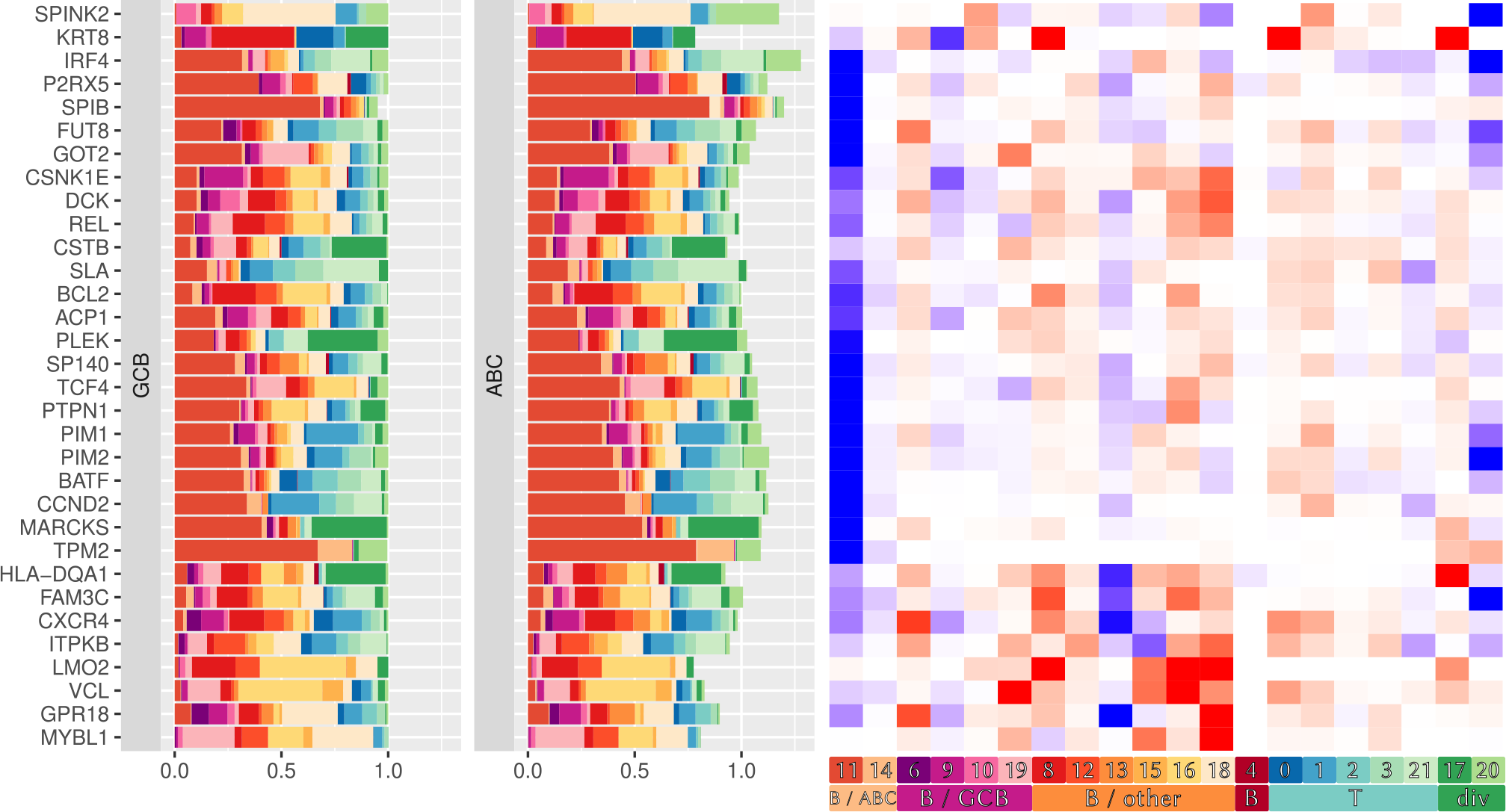
Volcano plot of cell type specific gene expression in cluster 17. Genes with absolute fold change higher than 0.5 and false discovery rate less than 0.05 appear in red. Among those we additionally labeled genes which belong to the stromal signatures.

**Fig. 6.**
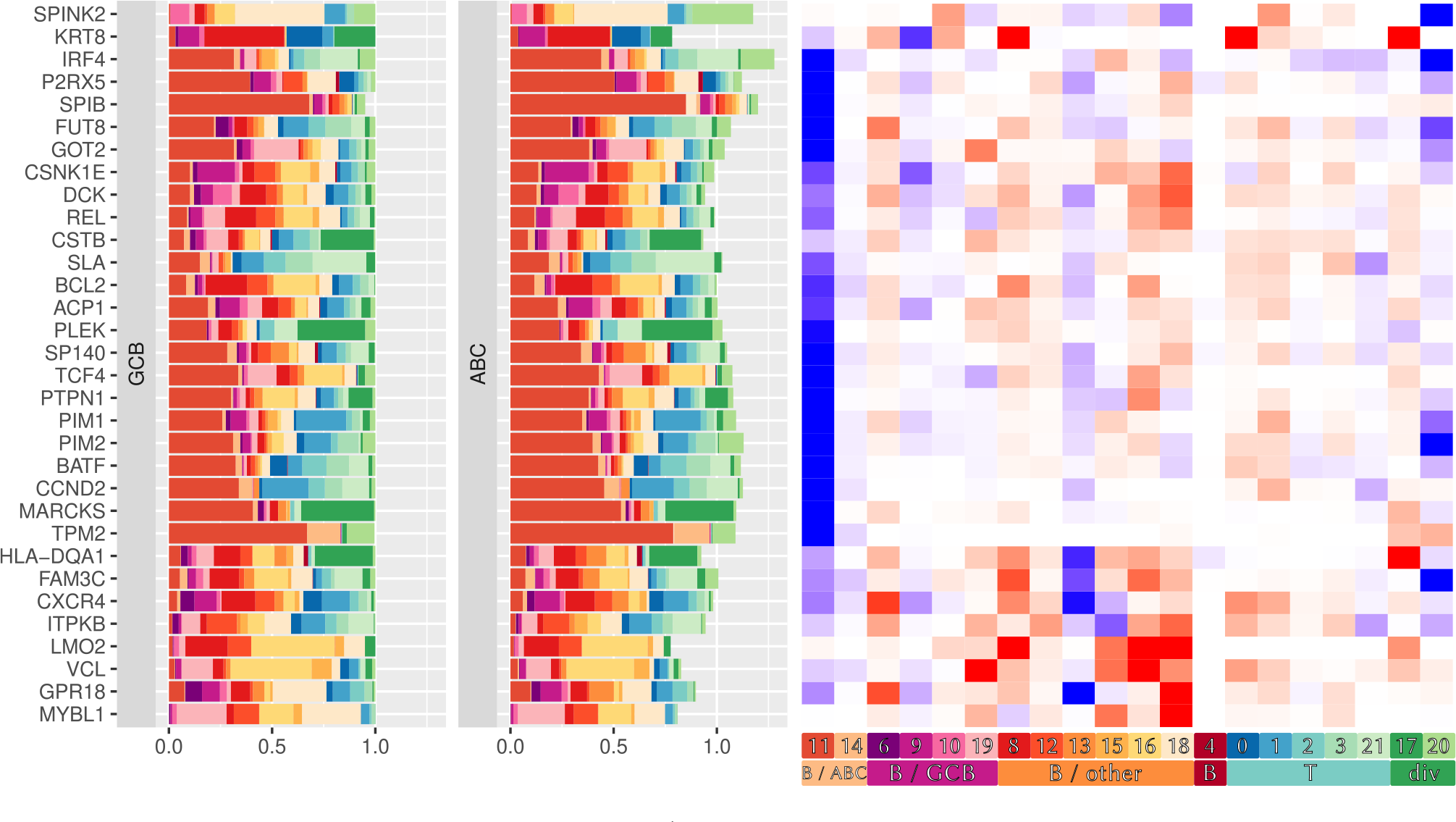
Cell type-specific expression of ABC/GCB signature genes from virtual tissues generated by tissueResolver: The bar plots on the left break down the total expression of a gene into contributions from various cell clusters, indicated by colors and averaged over the ABC and GCB groups. Due to differential expression between ABC and GCB lymphomas, these contributions do not add up to the same numbers. For better visualization, we normalized the GCB contributions to a constant, while the contributions in ABC lymphomas can add up to a higher or lower total amount, depending on the total expression of this gene in this group. The heatmap on the right highlights which cell populations (columns) are responsible for the differential expression of a signature gene (row). Blue indicates increased cell type-specific expression in ABC, whereas red means reduced expression in ABC than in GCB.

**Fig. 7.**
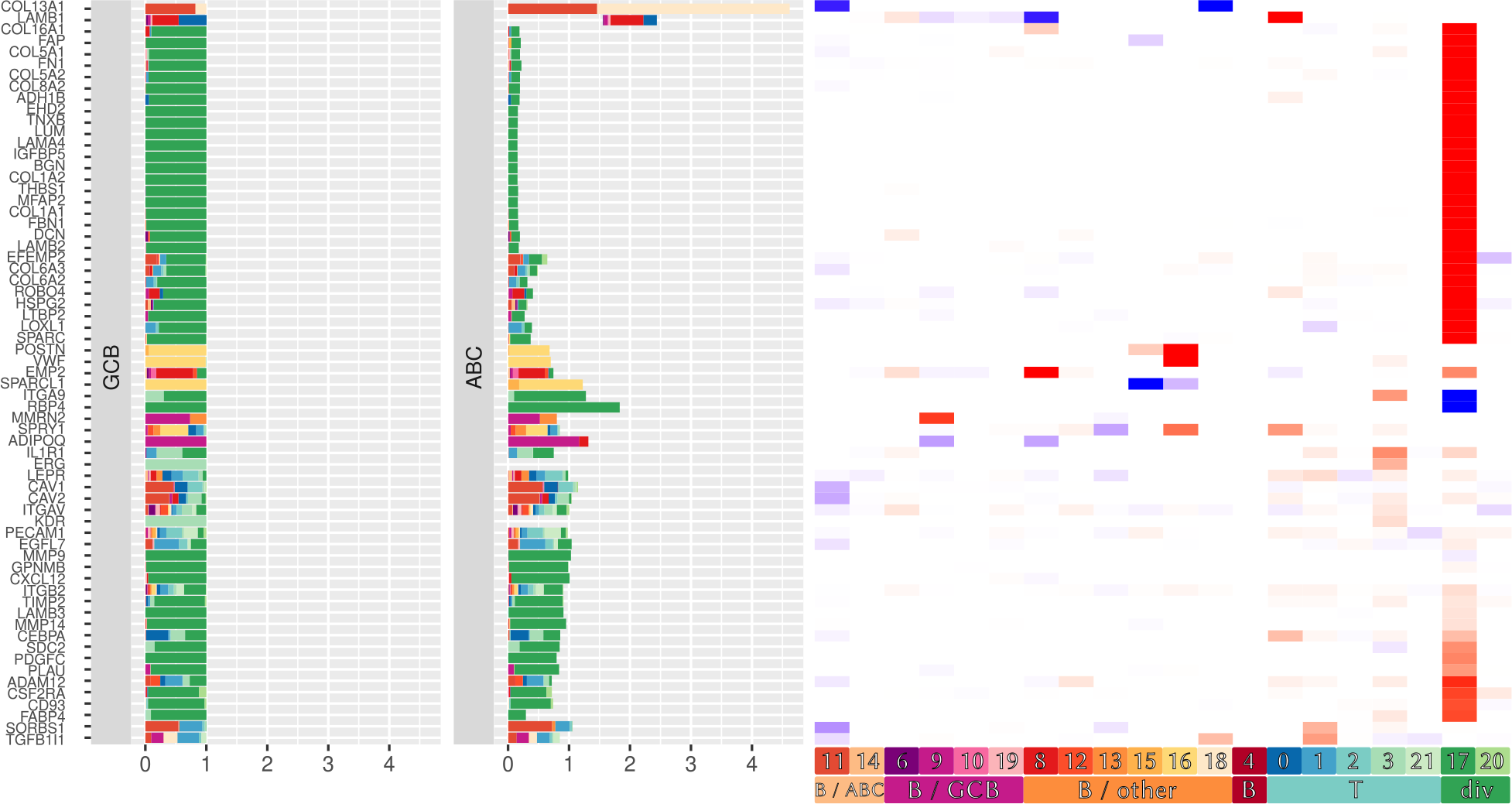
Cell type-specific expression of stromal signature genes. The Figure is organized analogously to Figure 6 above.

## IMPLEMENTATION

TissueResolver is implemented as an R package. To compute virtual tissues, it utilizes the L-BFGS-B algorithm [26], interfaced through the standard R package `optim` [27]. This algorithm is capable of handling the box constraints imposed in equation (1). Some interfaces use tibbles from the tidyverse library [28] for efficient data frame implementations. The tissueResolver package is available at https://github.com/spang-lab/tissueResolver and examples, simulation code and vignettes can be found at https://github.com/spang-lab/tissueResolver-docs.

## DISCUSSION

We introduced tissueResolver, a novel approach for estimating gene type-specific expression from bulk expression profiles. The method is entirely deconvolution-based, eliminating the need for a priori cell type definitions, thereby enhancing flexibility in its application compared to competing methods.

Not defining cell types upfront is akin to the approach presented in [13], with tissueResolver extending these ideas into a comprehensive, parameter-free deconvolution model. Similar considerations apply to BayesPrism [14]. As a complete Bayesian model, BayesPrism distinguishes itself from other methods by offering sound uncertainty quantifications for all its predictions. We see its downside in its reduced performance in our benchmark analysis and in its need to specify cell-types up-front.

Reconstructing virtual tissues is inherently constrained by the quality of the input data. Specifically, single-cell libraries must encompass cells similar to those in bulk tissues, and technical disparities between single-cell and bulk RNA-seq data should not hinder the reconstruction of the majority of genes. We defined quality metrics that allow to judge fits and interpretability of individual genes.

In summary, tissueResolver emerges as a novel player in cell type-specific gene expression reconstruction. It exhibits flexibility in its application and has shown outstanding performance in benchmark evaluations. TissueResolver enriches our understanding of tumor tissues, encompassing their micro-environment. While single-cell sequencing holds the promise of future cell-type-specific predictive gene expression signatures, tissueResolver facilitates this research in the present by unveiling concealed information within extensive existing datasets of bulk gene expression data.

## AUTHOR CONTRIBUTIONS STATEMENT

JS devised the algorithm, conducted simulations and data analysis, and authored the manuscript. PH contributed to data analysis, reviewed the implementation, and co-wrote the manuscript. MS aided in developing the R package and supported data preprocessing. ZN and MA provided valuable ideas and suggestions. TS, MH, and JS collaborated on developing the data management software utilized in the analysis. RS initiated and oversaw the project and co-wrote the manuscript.

## ACKNOWLEDGMENTS

The authors are grateful to Michael Rehli for carefully reading and improving the manuscript. This work is supported by BMBF (grant no. 031L0173). MA acknowledges funding by the Deutsche Forschungsgemeinschaft (DFG, German Research Foundation) [AL 2355/1-1].

## SUPPLEMENTAL MATERIAL

### Gene filtering

Before running tissueResolver, we reduced the number of genes to a common set of 1000 genes of interest to reduce runtime and restrict the optimization to a set of relevant genes. Since we are interested in the interpretation of gene signatures, all available genes from the ABC/GCB signature [18, 23] and the stromal signatures [25] were explicitly included in the final list of genes, irrespective of their expression or variances. The list was then filled up with top variable genes that were determined in the following way: First, ribosomal genes and all genes that are only available in either the bulk or the single cell datasets were removed. Then, the remaining genes were ranked separately by their bulk and single cell variances and the two lists were merged into a single list of joint, highly variable genes, alternating between genes from the single cell and bulk list.

Finally, we used edgeR [29, 30] to compute TMM normalization factors for the bulk on this subset of 1000 genes.

### Clustering of single cells

Although single cell datasets provided cell-type annotations based on a single cell analysis incorporating prior knowledge, we re-clustered the combined single cell datasets to analyse our computed virtual tissues and compute cell-type specific quantities. For the clustering, we followed the classical Seurat [31] pipeline and used Louvain clustering [32] on the top 15 PCA components with a resolution of 0.5, yielding 22 clusters, see table 1 and the UMAP embedding, fig. 3, for the relation to the original cell type annotations.

### Differentially expressed genes in virtual tissues

In the main text, we used tissueResolver to give meaning to existing signatures. In this section, we demonstrate that virtual tissues are also useful to explain *why* genes are differentially expressed, i.e. to understand what cell communities are responsible for an observed fold change.

First, we generate a list of differentially expressed genes by using edgeR [29, 30]: We use the existing ABC and GCB labels from the Schmitz dataset [19], (re-)compute normalization factors and estimate the dispersion. Since we consider only two groups, we use the classical exact test to compute *p*-values and false discovery rates. In the following, we restrict ourselves to genes with an absolute fold change greater than 2^0.8^ and a false discovery rate below 5 %. Next, we use our virtual tissue to compute cell-type specific gene expression values for the clusters defined above, cf. 1, and select genes that satisfy our quality criteria of *g*_*g*_ ≤ 0.5 and *v*_*g*_ ≤ 0.015, cf. fig. 9 and the discussion in the following section. Cell-type specific expression changes can be read off from fig. 8. We see that this reveals analogous regulatory effects as considering the classical ABC/GCB signature genes, see Section 4, with some overlap in the set of genes that were selected based on the fold change and quality criteria with those from the signature, e.g. BATF, PIM1 and PIM2. Again, the differences in ABC to GCB are largely attributed to the tumor itself. However, changes in clusters 17 and 20 are also clearly visible, e.g. for HSP90B1, HCK and CD99.

**Fig. 8.**
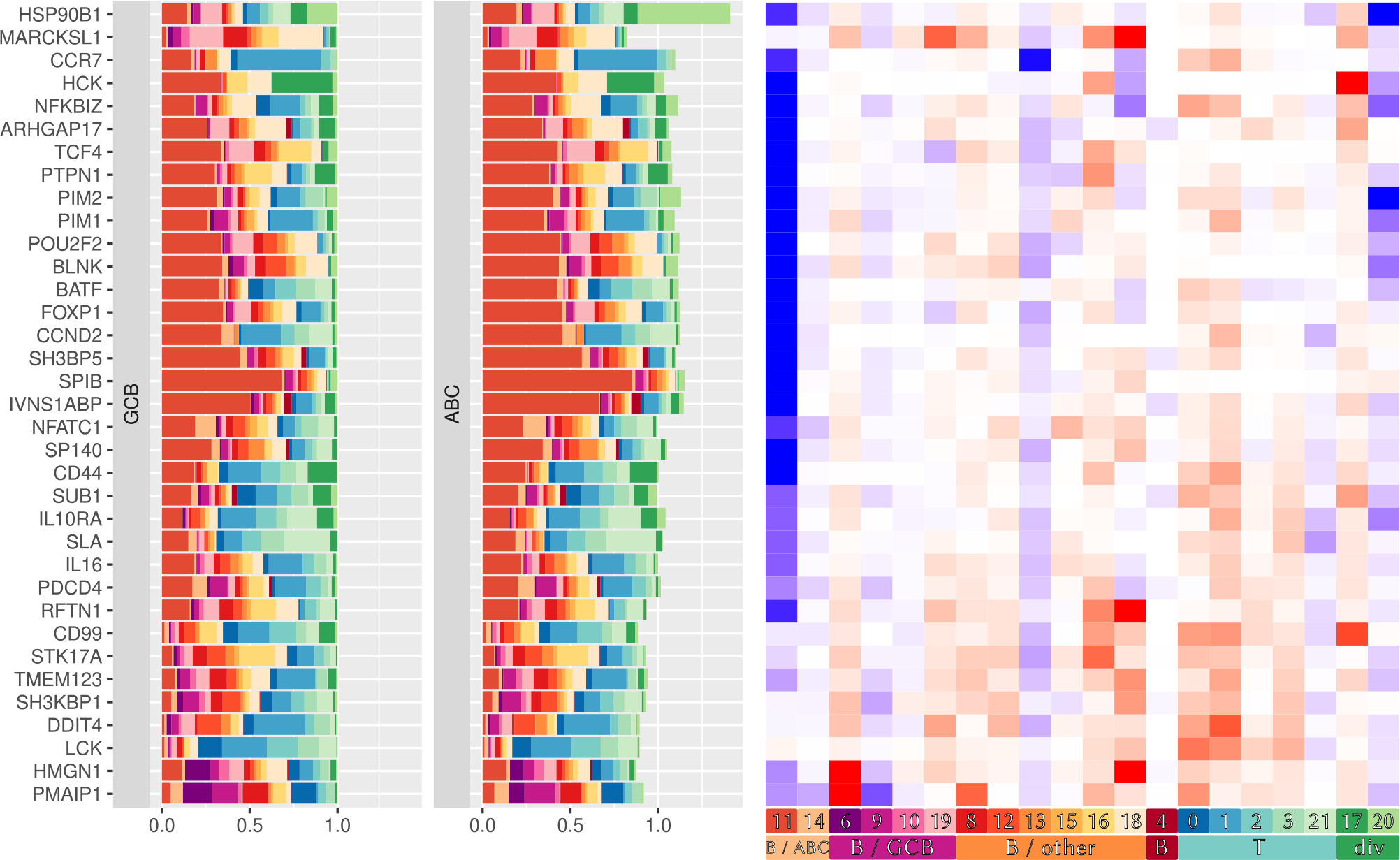
Genes with absolute log fold change *>* 0.8, FDR ≥ 0.05, a relative residual *g*_*g*_ ≥ 0.5, and an average variance of the relative residual *v*_*g*_ ≤ 0.015.

**Fig. 9.**
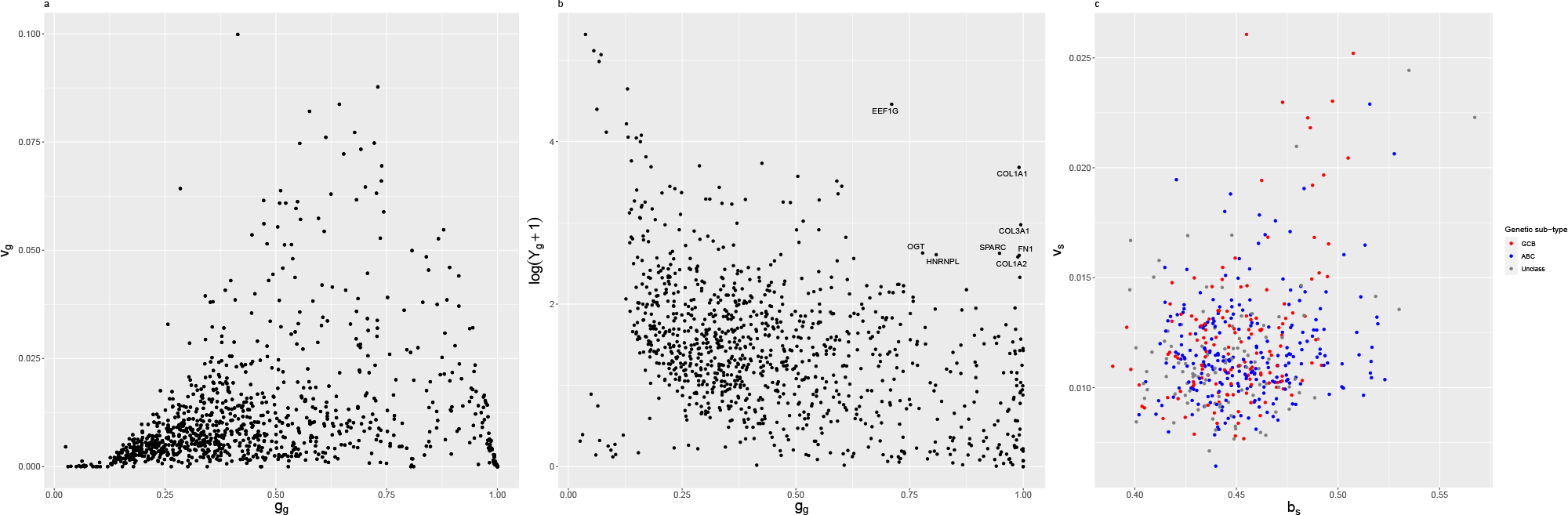
a) Mean relative residuals, eq. (5), for every gene *g*_*g*_ and their average bootstrap variance, 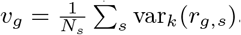. High variance values may indicate missing cell types in the reference. b) Mean relative residuals *g*_*g*_ and average expression 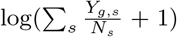 of genes. Highly expressed genes are typically well covered by the fit and display small relative residuals, whereas many of the lowly expressed genes are below the noise level and can hardly be interpreted. Genes that are highly expressed but show a large deviation indicate expression in cells that are not present in the single cell library. c)Mean relative residuals for every bulk *b*_*s*_ and their average bootstrap variance, 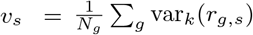. High variance values may indicate bulk tissues comprised of cells that are missing in the reference. We additionally colored each sample according to its genetic subtype. The equal distribution of colors indicates that there are no subtype specific differences in the quality of the fit.

### Quality control of virtual tissues

When screening for genes, it is crucial to judge the quality of virtual tissues and the trustworthiness of results on a per-gene basis. We do so now by interpreting the quality scores introduced in “Quality scores and gene selection”. From that section, we recall the gene-specific quality score *g*_*g*_ and the bulk-specific score *b*_*s*_, cf. eq. (5). Comparing the mean relative residuals of every gene *g*_*g*_ to their respective mean variances 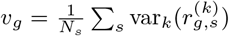 provides insight to how complete genes can be explained with the available single cell library, see Figure 9 a): Genes with small residuals that are consistently explained across bootstrap samples (small variance) are fully represented by the single cell reference. Genes with an increased residual are often also less stable. This is an indication that cell identities are missing and the gene expression values in the virtual tissue often deviate from their bulk values. Genes that could not be fit at all (*g*_*g*_ ≈ 1) show smaller values of *v*_*g*_, because no cells were available that could explain their bulk expression. For these reasons, Figure 8 displays only genes that fulfil *g*_*g*_ ≤ 0.5 and *v*_*g*_ ≤ 0.015 and that are differential in the bulks.

Unsurprisingly, genes that show the smallest relative residuals are also the ones with the highest gene expression in the bulk, cf. fig. 9 b), because they contribute the most to the loss function, eq. (1). These genes that are highly expressed and can be fitted well serve as driver genes to tissueResolver and help in selecting important cells, pinning down their specific weights, whereas cells that do not contribute enough to the loss or co-express genes that would increase the total loss are excluded by assigning small weights. For genes with high relative residuals, in contrast, we typically also observe low bulk expression, with some exceptions, namely genes that are not well covered by the single cell library. Possible reasons for a poor performance in highly expressed genes may also include technological differences between the single cell and bulk RNA-seq workflows.

Lastly, there may also be cases where only a subset of bulks can be fitted sufficiently well. Complete quality control therefore also demands for an assessment of the scores at the bulk level. We considered 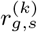 averaged now over genes, *b*_*s*_, and compared the bulk-specific score *b*_*s*_ to the average bootstrap variance 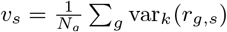, see Figure 9 c). In our specific case, the majority of bulks has been fitted with good or acceptable residuals and small variances. If, however, we would observe high variances, these instable bulks contained significant contributions from cells that are not in our single cell library. By looking at the annotated subtype [19], we deduce that in our case this is not the case and the quality of the fit is almost independent of the tissue’s genetic signature and so we do not miss any important phenotypes in our library.

In summary, we see that virtual tissues computed by tissueResolver are capable of explaining bulk expression as long as the used single cell library contains a reasonable amount of cells essentially covering the heterogeneity of the bulk, e.g. the groups that are compared should also be represented in the single cell data. For the cells of origin of DLBCL constituting the ABC/GCB signature, namely the B-cells, this is the case by observing several patient specific B-cell clusters, see Figure 3 and Table 1. We stress that these are different patients than those in the bulk datasets and yet tissueResolver chose a combination of cells from only 20 patients that resembles the bulk expression in a way that it recovers the genetic subtype of the tumors. When screening for genes, however, some caution is required as typically, there exist genes that cannot be fully explained by the available single cells. In the following, we will demonstrate that some information on these genes may still be amenable for interpretation.

Let us reconsider the genes in the stromal signature, cf. Figure 7. We show the quality scores for these genes in Figure 10.1 a) and b). First of all, we see that many of these genes are only moderately expressed in the bulk and show a high relative residual, i.e., using the available single cell data, the virtual tissues could not fully explain the entire gene expression of the bulks. However, we observe, that many of these genes are attributed to cluster 17 and behave stable across all bootstrap runs (small *v*_*g*_). Also, cluster 17 is assigned stable weights across the two groups and the total sum of weights changes only slightly between them, cf. fig. 4. In other words: The cell weights in cluster 17 that are chosen to explain the bulks are pinned down precisely by highly expressed driver genes, cf. also fig. 8 where cluster 17 contributes significantly to highly expressed genes with a large foldchange and good quality scores. One of the key features of tissueResolver is that the reference matrix consists of true, actual cells. Therefore, although the bulk expression is not fully explained (large *g*_*g*_), the *contribution* of cluster 17 to the total expression can be trusted because in that cluster these genes are co-expressed with genes that allow to predict the frequencies and weights of the relevant cell population. The integrity of the cells’ profiles allow us to view the resulting tissue as a predictor also for the lowly expressed genes of the stromal signature. This is only possible as our algorithm is not demanding for a priori cell clustering, thus, giving each single cell an unbiased chance to show its relevance in differentiating between genetic signatures.

**Fig. 10.**
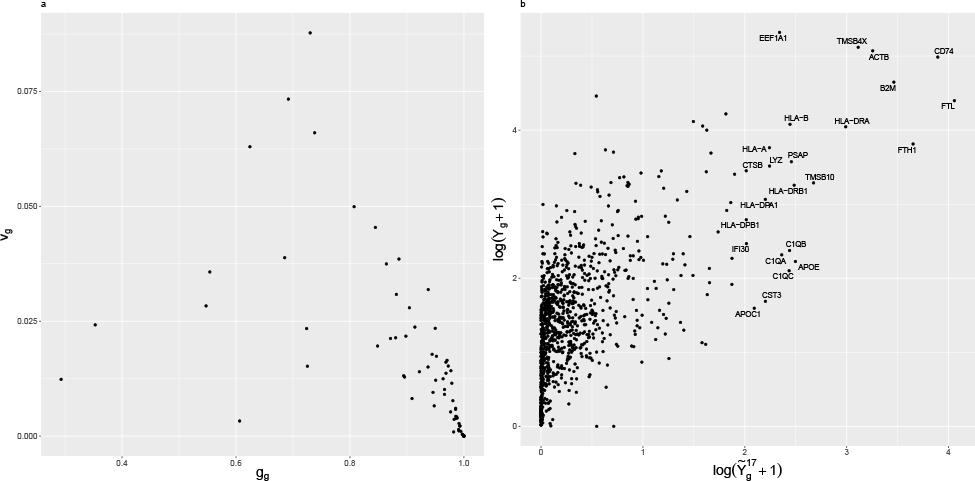
Quality scores for stromal genes. See fig. 9 for details. b) shows the mean expression of each gene declared by cluster 17. We see that there are genes which are highly expressed both in cluster 17 and the actual bulk.

With this in mind, we are tempted to have a closer look at individual clusters that are consistently integrated into virtual tissues. In fig. 5, we show fold changes and corresponding false discovery rate between the two conditions in a classical volcano plot, but only using the specific regulation found in cluster 17. Comparing the differential cell-type specific regulatory genes with those in the stromal signature (labels in the figure), we find that the signature is well recovered by these cells and, had we screened for a cell-type specific signature in our data, we would likely have ended up with a similar result as Lenz et al. [25]. This highlights the use of tissueResolver as a tool to identify cellular mechanisms and signatures that would otherwise remain hidden in the bulks.

### Simulation details

In the following we provide our test setup as pseudo code, Algorithm 1, providing additional details and a step by step pipeline supplementing the explanations in Section “Simulations”. Note that for BayesPrism, we included two possible scenarios, one where we use exclusively the annotated cell-types from the single cell dataset (“bp-nosub”, step 4b) and one where we subcluster every annotated cell-type (“bp-sub”, step 4c, also see section “Clustering of single cells” for details on the clustering) and provide these as “cellstates”. We perform this cluster step in every iteration, i.e. whenever the single cell library changes. The simulation is repeated five times to determine averages and standard errors of fig. 2 (step 5). Runnable R code for both the BayesPrism as well as the tissueResolver benchmarks is provided along with the vignettes on https://github.com/spang-lab/tissueResolver-docs.

The selection of genes that were modified, G_mod_, was done in the following way: From all annotated CD8+ T-cells, we determined the set of genes that showed non-zero expression in at least 60 % of all these cells and further required that the genes are also expressed in 60 % of all available cells (across cell types). This avoids the selection of highly specific genes that would result in a trivial deconvolution task, as the genes differentiating between the two bulk groups were then only expressed in the cell type we have modified and acted as marker genes.

#### Algorithm 1

Simulations for benchmarking tissueResolver and BayesPrism

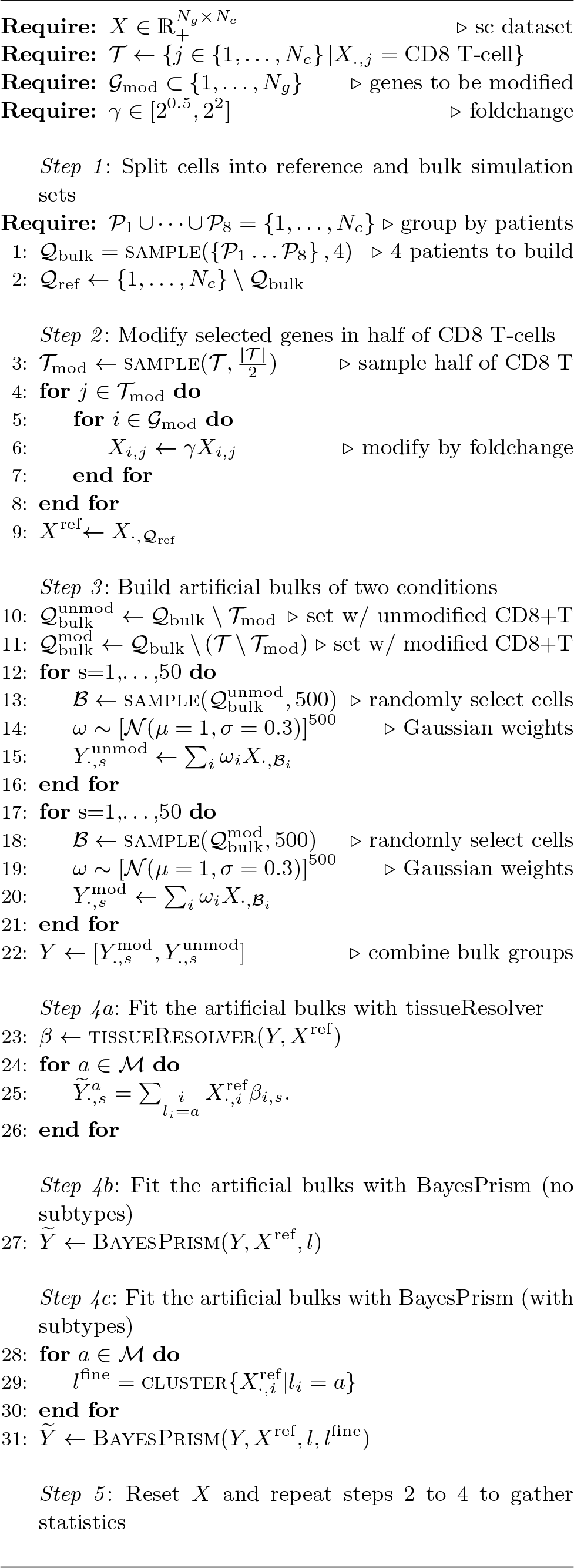

## Footnotes

1 The data of [17] is publicly available under GEO accession numbers GSE182436 and GSE182434

2 The data of [19] is publicly available in GDC.

3 The scRNA-seq count data of [20] is publicly available under heiDATA ID VRJUNV and EGA accession number EGAS00001004335.

